# Protocol combining RNA interference and regeneration assays in planarian embryos

**DOI:** 10.1101/2025.07.21.666018

**Authors:** Ennis W. Deihl, Clare L. T. Booth, Erin L. Davies

## Abstract

Planarian flatworms gradually acquire whole-body regeneration abilities during late embryonic and juvenile development. Here we show how to perturb gene function with RNA interference (RNAi) in *S. polychroa* embryos at developmental stages capable of wound healing following amputation. We describe embryo staging, embryo amputation and double-stranded RNA soaking, phenotype analysis, and techniques for qualitative and quantitative assessment of RNAi knock-down efficacy. This protocol is adaptable for use with embryonic stages amenable to *ex vivo* culturing and with other embryo-producing flatworm species. For complete details on the use and execution of this protocol, please refer to ^1^.

**Graphical abstract:** 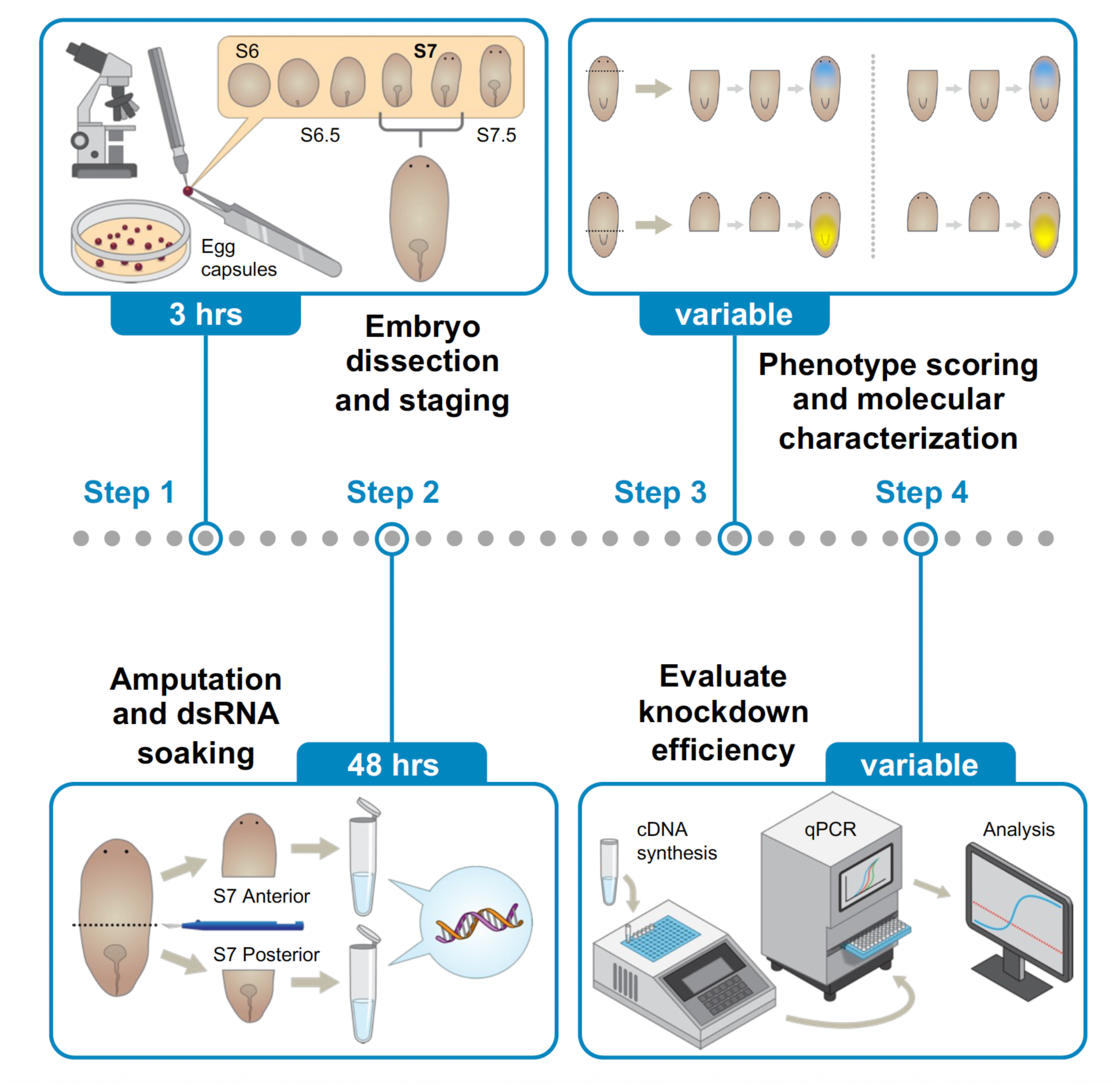

## Before you begin

RNAi knock-downs enable interrogation of loss-of-function phenotypes in animal models for which forward genetics and transgenic approaches are not readily available, including highly regenerative planarian flatworms. For the past two decades, RNAi screens have shed light on genetic regulatory networks implicated in adult pluripotent stem cell self-renewal^2,3^, cell fate specification and tissue-specific differentiation programs^4–9^, reproduction^10–12^, and regeneration^13–15^. These assays are typically performed by introducing double stranded (ds)RNA for gene(s) of interest into the gut by feeding or injection^16–18^, though dsRNA soaking has been reported for adult regenerating fragments^19^. In contrast, little progress has been made in the adaptation of RNAi protocols for use in developing planarian embryos^20^.

This protocol explains how to conduct dsRNA soaking-mediated RNAi knock-downs in wound healing-competent Stage 7 (S7) *S. polychroa* embryos following amputation^1^ (**Figure 1A**). Experimenters should have access to a *S. polychroa* breeding colony, maintained in 1x Montjuic water^21^ as described in^1^, with egg capsules collected daily and stored in an unhumidified 20°C incubator. This protocol may be adaptable for use with other embryo-producing planarians and potentially other aquatic organisms.

**Figure 1:**
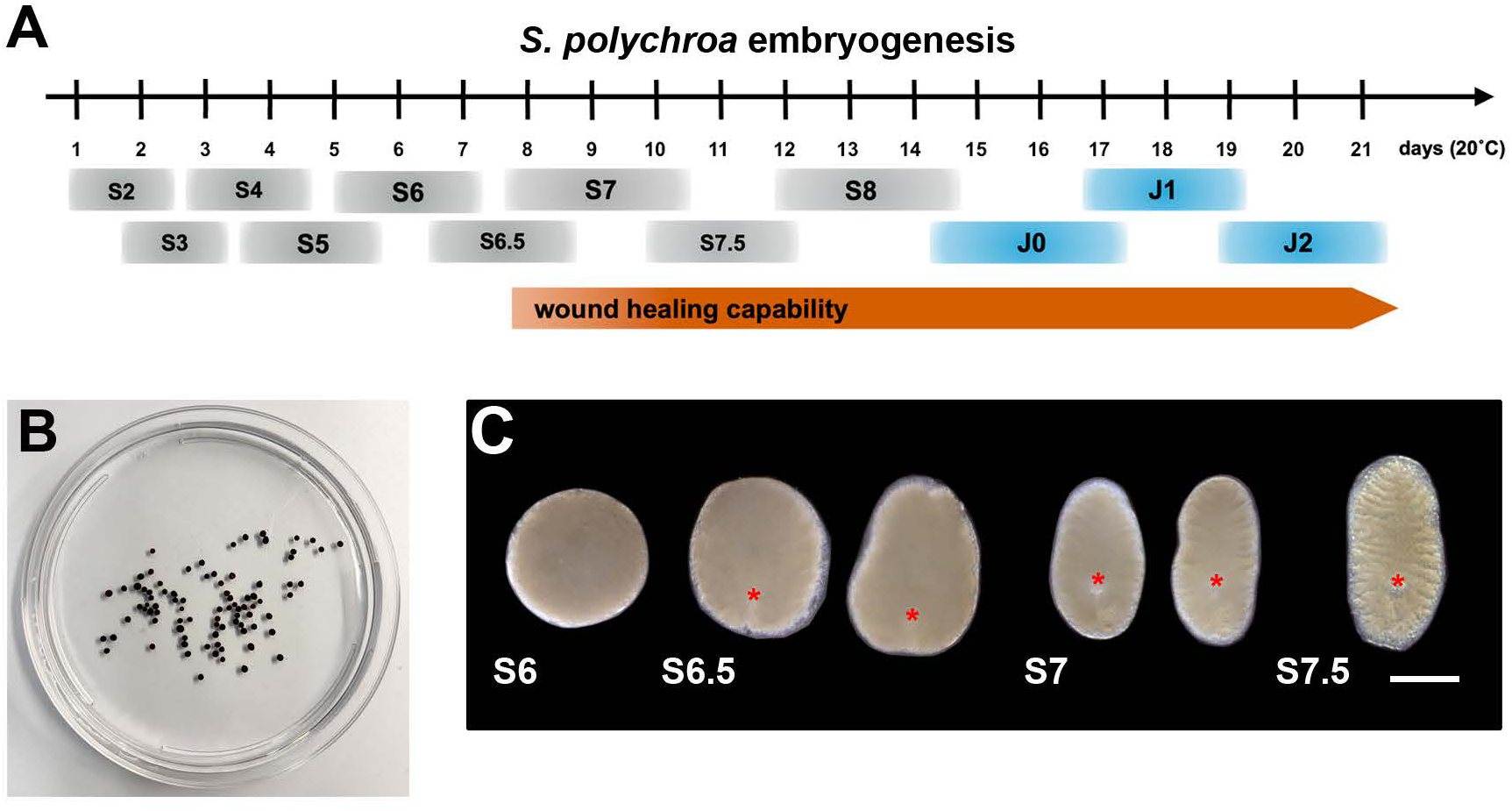
*S. polychroa* embryos are staged using chronological, morphological, and behavioral criteria. 1A. *S. polychroa* embryonic development proceeds through Stages (S) 1-8. Hatching marks Juvenile 0 (J0). Embryos are capable of surviving amputations from S7 onward. 1B. Quinine-tanned *S. polychroa* egg capsules. Gravid egg capsules may contain one or several embryos. 1C. Live images, *S. polychroa* embryos, S6 – S7.5. S6.5, S7, S7.5: Anterior up. S6.5, S7: ventral views. S7.5: dorsal view. Asterisks: definitive pharynx. Scale bar: 500 µm.

The experimenter must generate dsRNAs for target gene(s) prior to starting our protocol. We recommend making dsRNA for a positive control, i.e., a gene that produces a penetrant, morphologically striking phenotype upon knock-down, like *Spol-ß-catenin-1*^1^. dsRNA for a sequence that is absent from *S. polychroa* (or your research organism of interest) should be generated for use as a negative control. *C. elegans unc-22*^22^ or GFP dsRNA are frequently used as negative controls for planarian RNAi experiments. One or more non-overlapping dsRNAs, each approximately 500 bp in length, should be generated complementary to your gene of interest. We recommend subcloning sequence for your target into a vector containing two T7 RNA polymerase binding sites in opposite orientation flanking the insertion site (e.g., pJC53.2 Addgene #26536^23^ or pDL1 Addgene #182263^24^) and verifying clone identity by whole plasmid sequencing. dsRNA can be synthesized *in vitro* from purified PCR templates, with 5’ T7 RNA polymerase binding sites flanking the amplicon of interest, as described in^18,25^. Alternatively, vectors can be transformed into HT115 bacteria for dsRNA production using IPTG-inducible expression of T7 RNA polymerase, followed by dsRNA purification as described in ^26^.

We explain steps required for successful dissection and staging of *S. polychroa* embryos, S7 embryo amputation, and dsRNA soaking of S7 cut fragments. We provide examples and advice on scoring and characterizing RNAi knock-down phenotypes, as well as qualitative and quantitative measures of RNAi efficacy.

### Institutional permissions (if applicable)

All experiments should be conducted in accordance with applicable institutional and national guidelines and safety regulations.

### Prepare solutions

**Timing: [1-2 hours]**

The experimenter should prepare stock solutions using recipes provided in the Materials and Equipment Set Up section.

1. Prepare 50 mg/mL gentamicin sulfate solution.

a. Dissolve 10 g gentamicin sulfate powder in deionized water, adjusting final volume to 200 mL.
b. Filter sterilize using a disposable bottle top filter (0.2 µm membrane).
c. Store at 4°C.
2. Prepare 5x Holfreter’s solution.

a. Dissolve sodium chloride, sodium bicarbonate, potassium chloride, magnesium sulfate, calcium chloride dihydrate, and dextrose in quantities indicated in deionized water.
b. Adjust to pH 7.5, using 1 N HCl and 1 N NaOH as necessary.
c. Add deionized water to achieve a final volume of 200 mL.
d. Filter sterilize using a disposable bottle top filter (0.2 µm membrane).
e. Store at room temperature.
3. Prepare 1x Holfreter’s solution.

a. Dissolve sodium chloride, sodium bicarbonate, potassium chloride, magnesium sulfate, calcium chloride dihydrate, and dextrose in quantities indicated in deionized water.
b. Adjust to pH 7.5, using 1 N HCl and 1 N NaOH as necessary.
c. Add deionized water to achieve a final volume of 1 L.
d. Filter sterilize using a disposable bottle top filter (0.2 µm membrane).
e. Store at room temperature.
4. Prepare 1x Holfreter’s+ 100 µg/mL gentamicin solution.

a. Add 1 mL of 50 mg/mL gentamicin sulfate stock solution to 500 mL 1x Holfreter’s solution. Store at room temperature and use within 2 weeks.
5. Prepare 10% bleach solution.

a. Pipet 10 mL bleach into a polypropylene squirt bottle containing 90 mL deionized water. **Note:** Wear gloves, protective eyewear, and a lab coat when handling bleach.
6. Prepare 1% low melt agarose coated petri dishes.

a. Dissolve 0.5 g low melt agarose in 50mL of pre-warmed 1x Holfreter’s solution. We recommend using a microwave-safe Erlenmeyer flask and short (10-20 second) pulses with repeated, gentle swirling until the agarose is completely dissolved. Adjust final volume to 50 mL to account for any loss due to evaporation. **Note:** Wear gloves, protective eyewear, and a lab coat when handling molten agarose. Handle the hot flask using an insulated potholder. Be mindful that the agarose solution may bubble up or boil over easily.
b. Using a cut p1000 tip, apply 1.0 mL melted agarose per 35 mm petri dish. Swirl gently to ensure the agarose evenly coats the bottom of the dish, then allow agarose to solidify at room temperature.
c. Cap and store agarose-coated plates in a sealed Tupperware container at 4°C for up to 2 weeks.

### Embryo Dissection and Staging

**Timing: [variable]**

7. Bleach intact egg capsules to minimize microbial and protist contamination.

a. Transfer *S. polychroa* egg capsules with a disposable #691 transfer pipet into a clean 100 mm petri dish. Remove untanned (yellow) egg capsules, broken egg capsules, and debris (**Figure 1B**). **Note:** We recommend bleaching egg capsules from daily collections in separate dated dishes. Dissect 8-9 day post-egg capsule deposition (dped) egg capsules to find Stage 7 (S7) embryos. The date the egg capsules are collected is 1 dped (provided egg capsules are collected daily).
b. Remove fluid and submerge egg capsules in 20 mL 10% bleach for 3 minutes.
c. Rinse egg capsules thoroughly with at least 4 quick exchanges of 1x Holfreter’s solution. **CRITICAL**: Repeated washes will guard against bleach carryover.
d. Resuspend egg capsules in 20 mL 1x Holfreter’s + 100 µg/mL gentamicin sulfate.
8. Dissect embryos from egg capsules.

a. Insert a new #000 insect pin into the pin holder. Spray down pin and forceps with 70% ethanol prior to use and wipe dry with a kimwipe.
b. Gently hold an egg capsule with the forceps and immobilize it against the dish bottom. Poke the egg capsule gently with the insect pin, puncturing or cracking the shell.
c. Use the pin and forceps to cleave and open the egg capsule, allowing embryos and yolk to spill out into the culture media. Carefully separate embryos from the yolk and transfer embryos, using a cut p200 pipet tip, to a 1% low melt agarose-coated 33 mm petri dish containing 1x Holfreter’s media + 100 µg/mL gentamicin. Discard injured embryos. **Note:** Gravid egg capsules frequently contain more than one embryo. Embryos within an egg capsule may vary slightly in their size, morphology, and developmental stage. **Note**: We recommend transferring embryos to petri dishes seeded with 1% low melt agarose beds to minimize embryo sticking and injury arising from direct contact with the polystyrene dish bottom.
d. Stage embryos and divide them into experimental groups (**See Figure 1C, Troubleshooting Problem 1**^1,27^.

i. **Stage 6 (S6) embryos** (age range: 6 – 8 dped): opaque, yolk-filled spheres approximately 500 µm or greater in diameter. Embryos are immotile (**Figure 1C)**.
ii. **Stage 6.5 (S6.5) embryos** (age range: 7-9 dped): opaque, yolk-filled embryos undergoing elongation along the main body axis. Morphology varies; embryos may be ovoid or pyramidal. Flattening begins along the dorsoventral axis. The definitive pharynx primordium and developing tail stripe are visible in the posterior third of the embryo. Embryos are immotile (**Figure 1C).**
iii. **Stage 7 (S7) embryos** (age range: 8-10 dped). Embryos are elongated, with a defined head and tail. The melanized optic cup cells of the eyes first become visible on the dorsal side of the head. The triclad gut begins taking shape, with primary and secondary branches becoming visible; branching may be more pronounced in the anterior relative to the posterior. The definitive pharynx primordium is visible in the posterior half of the embryo. The tail stripe is visible and divides the posterior primary gut branches from one another. Pigment cells are sparse but sometimes present on the dorsoanterior side **(Figure 1C)**. Onset of gliding locomotion. When embryos are positioned ventral side up, they cannot curl their heads toward the ventral surface, nor can they twist to right themselves.
iv. **Stage 7.5 (S7.5) embryos** (age range: 9-11 dped). Eyes are small but visible. Gut branches are better resolved relative to S7, and branching is visible along more of the AP axis. A clear zone of tissue, superficial to the gut, extends around the animal (**Figure 1C).** When embryos are positioned ventral side up, they can curl their heads toward the ventral surface and can twist to right themselves.

### Embryo amputation and dsRNA-soaking of embryo fragments

**Timing: [1 hour; 2 days]**

9. Prepare dsRNA soaking solutions for positive control, negative control, and experimental knock-downs. We recommend preparing 150 µl solution in 0.2 mL snap cap PCR tubes or 0.5 mL microfuge tubes, at room temperature, prior to starting amputations. Up to 25 fragments can be soaked in 150 µl dsRNA-containing solution.

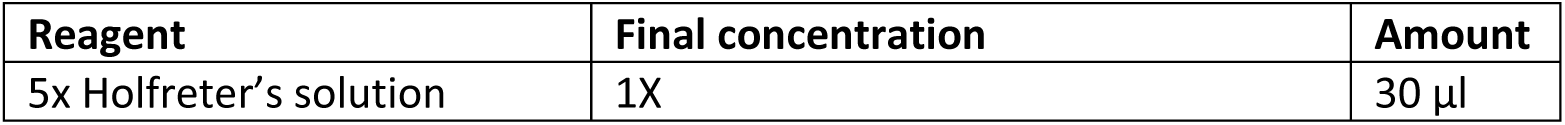
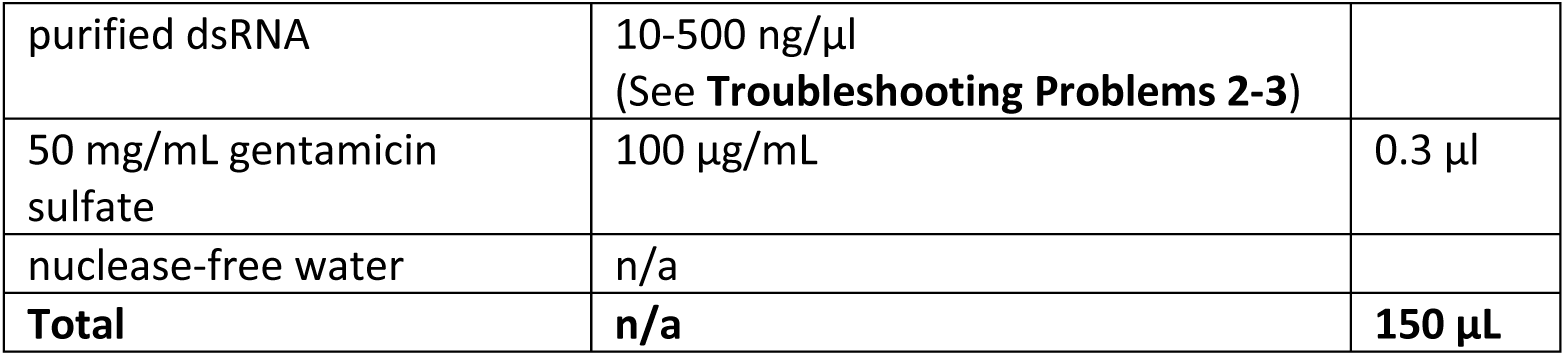 **Note:** We recommend allotting approximately 20-25 S7 embryo fragments per dsRNA dose tested. Volumes and fragment numbers can be scaled up as needed for downstream validation and phenotypic characterization. **Note:** We advise against mixing amputated fragments of different types within the same tube, even when both fragments are to be soaked in the same dsRNA and dose. Create separate dsRNA solutions for each fragment type, gene, and dose assayed.
10. For each dsRNA and dose assayed, select approximately 20-25 intact S7 embryos for amputation.

a. Transfer embryos to a dish lid containing 1x Holfreter’s + 100 µg/mL gentamicin with a cut p200 pipet. Do not expose embryos to air. **CRITICAL**: Proper staging is critical for experimental reproducibility and will impact the survival and regenerative ability of cut embryo fragments (See **Troubleshooting Problem 1)**. We recommend performing transverse amputations at S7 or later. The vast majority of S7 anterior and posterior fragments, produced by making a single cut immediately anterior to the pharynx, undergo wound closure and survive through 14 days post-cut (dpc)^1^. In contrast, less than 50% of S6.5 anterior and posterior fragments undergo successful wound closure^1^. **CRITICAL**: Embryos exhibit developmental stage- and axial position-dependent differences in their regenerative abilities prior to hatching^1^. Experimenters must carefully consider fragment type, axial position of the cut plane, wound orientation, and developmental stage when designing knock-down experiments (see **Troubleshooting Problem 4-5**).
11. Amputate S7 embryos using 22.5° surgical micro knife and immediately transfer fragments to dsRNA solution using a cut low-retention p200 pipet tip.

a. Use the position of the developing eyes to locate the anterior head margin and the position of the pharynx primordium to locate the ventral posterior domain of the embryo prior to cutting (**Figure 1C, Troubleshooting tips 4-5**).
b. Apply minimal pressure needed to make the desired cut.
c. Use a cut p200 tip to transfer fragments to dsRNA solution. Embryo fragments will settle close to the tip edge and can be transferred with minimal fluid carryover to the dsRNA-containing solution by gravity when the tip meets the surface of the solution.
d. Between samples, wipe down the surgical blade with a 70% ethanol-soaked kimwipe to remove mucus and debris. **Note:** Avoid cross-contamination by replacing pipet tips between experimental samples. **Note:** If available, blunt cut low retention p200 tips are recommended for embryo transfers.
12. When amputations are complete, firmly cap tubes and lay tubes on their sides to allow the fragments to spread out relative to one another. Gently rolling the tube over will help fragments to spread out if they are clustering together. Incubate fragments in dsRNA solution in a non-humidified incubator at 20°C in the dark.
13. Following the 2-day incubation in dsRNA solution, fragments are transferred to 35 mm dishes containing fresh 1x Holfreter’s media + 100 µg/mL gentamicin sulfate.

a. Use a cut p200 tip for the fragment transfers. Flush out fragments, allowing for transfer of dsRNA solution to the new dish. Avoid exposing fragments to air during the transfer.
b. Remove debris, including yolk and lysed fragments, from the dish. **Note:** Avoid cross**-**contamination by replacing pipet tips between experimental samples. **Note:** Label dish side and lid with the start date, fragment stage and type, dsRNA name, and dose. **Note:** Avoid overfilling the 35mm dish with 1x Holfreter’s media as fragments can crawl onto the lid, potentially resulting in the loss of experimental samples.
14. Perform fluid exchanges at 3 days post-cut (3 dpc) and every 2-3 days thereafter. Do not expose fragments exposed to air when exchanging fluids.

a. At 3 dpc, replace fluid with 1x Holfreter’s media + 100 µg/mL gentamicin sulfate.
b. At 4-5 dpc, replace fluid with 1:1 (v/v) 1x Holfreter’s media: 1x Montjuic planarian water + 100 µg/mL gentamicin sulfate.
c. From 5 dpc – 14 dpc, perform fluid exchanges with 1x Montjuic planarian water + 100 µg/mL gentamicin sulfate.

### Vital phenotype scoring and imaging

**Timing: [variable]**

15. We recommend scoring fragments for visible morphological phenotypes at 2 dpc, 7 dpc, and 14 dpc using a dissecting stereomicroscope.

a. Score surviving fragments (n) at all three time points.
b. Note the number of 2 dpc fragments that underwent successful wound closure. **Note:** Most fragments will undergo complete wound closure by 2 dpc ^1^.
c. Score regeneration of missing tissue(s)/structures in negative control and experimental knock-down animals at 7 dpc and 14 dpc.

i. Anterior fragments: score for regeneration of posterior structures, including the pharynx and tail. **Note:** To assess regeneration quality, we recommend scoring pharynx function by performing feeding assays^9^ after 10 dpc. Tail morphology differences (e.g., incidence of forked tails) are also scorable.
ii. Posterior fragments: score for regeneration of anterior structures, including the head and eyes. **Note:** To assess regeneration ability and quality, we recommend scoring eye number (e.g., eyeless, 1 eye, 2 eyes, > 2 eyes) and eye patterning (e.g., incidence of cyclopic eyes). Head regeneration may also be assessed, in part, through behavior by assaying whether fragments undergo gliding locomotion.

**Note:** Many regeneration phenotypes are visible by 7 dpc; some will be easier to score at 14 dpc, when regenerated structures in controls are larger.

16. Capture representative images of phenotypes using a stereomicroscope fitted with a color camera.

a. Prior to imaging, transfer animals to clean petri dishes containing chilled 1x Montjuic water. Cold water will slow movement of the worms, making them easier to image. **Note:** We recommend storing a squirt bottle of 1x Montjuic at 4°C for on-demand use.
b. We recommend using a black background and lighting specimens from the top and/or sides using goosenecks.

### Optional: Harvesting knock-down animals in TRIzol for total RNA extraction

**Timing: [10 minutes for homogenization; 2 hours for total RNA extraction]**

Steps 17 and 18 are only required when the experimenters intend to run qRT-PCR to quantify knock-down efficacy.

17. Homogenize knock-down samples in TRIzol for total RNA extraction.

a. Transfer five representative 14 dpc fragments per sample to labeled 1.5 mL microfuge tubes using cut p200 tips.
b. Remove planarian water from samples using a #232 transfer pipet, taking care not to disturb the fragments. **CRITICAL:** We recommend performing steps 17c-e in a chemical fume hood while wearing gloves, a lab coat, and eye protection.
c. Immediately add 100 µl TRIzol to each tube.
d. Homogenize samples completely using a disposable plastic pestle and handheld tissue homogenizer, providing several 1-2 second pulses to each sample. Continue pulsing until fragments are no longer visible in the tube.

i. Use separate pestles for each sample to avoid cross-contamination. **Note:** If a tissue homogenizer is not available, disrupt tissue by manually grinding fragments using disposable pestles and/or repeated pipetting with a p200 until tissue is no longer visible.
e. Add 900 µl trizol to each sample, pipetting gently to mix, bringing sample volumes to V_total_= 1 mL. **Note:** Homogenized samples can be stored at −20°C or −80°C until total RNA extractions are performed.
18. Perform total RNA extractions following the protocol provided by the manufacturer (TRIzol, Invitrogen # 15596026).

a. Follow recommendations for working with small volumes of tissue, including the addition of 0.5 µl RNase-free glycogen prior to total RNA precipitation with isopropanol.
b. Resuspend total RNA in 20 µl nuclease free water.
c. Quantify and assess quality of total RNA using a nanodrop spectrophotometer or Agilent Bioanalyzer RNA Nano Chip.

**CRITICAL:** Perform total RNA extractions in a chemical fume hood while wearing PPE (lab coat, gloves, safety glasses). Follow hazardous waste collection and disposal guidelines for TRIzol, choloroform, isopropanol, and ethanol.

**Note:** Use total RNA to generate cDNA from *control(RNAi)* and *target gene(RNAi)* samples. We recommend using the Invitrogen Superscript III First Strand Synthesis Kit (#18080051), priming with oligodT or random hexamers, following the manufacturer’s instructions. Use cDNAs as templates in qRT-PCR reactions, as described in^1^.

### Optional: Fixing knock-down animals for whole mount *in situ* hybridization or immunostaining

Steps 19 – 26 are required to fix 14 dpc regenerates for further characterization of phenotypes with molecular markers.

**Timing: [2 hours]**

**Prepare solutions**

19. Prepare 5% (w/v) N-acetyl cysteine (NAC) in 1x PBS.

a. Dissolve 0.5 g NAC per 10 mL 1x PBS. Shake well, until fully dissolved. **Note:** We recommend preparing 5% NAC fresh to ensure effective removal of mucous prior to fixation.
20. Prepare 4% formaldehyde in 1x PBSTx (1x PBS +0.5 % Triton X-100). **CRITICAL:** Prepare fixative in a chemical fume hood while wearing PPE (lab coat, gloves, safety glasses). Follow hazardous waste collection guidelines for formaldehyde-containing solutions.
21. Perform anti-mucolytic treatment with 5% N-Acetyl Cysteine in 1x PBS.

a. Transfer regenerated 14 dpc animals to clean petri dishes, n ∼ 20 in a 60 x 15 mm petri dish, or up to n ∼ 50 into a 100 mm dish.
b. Working with one dish at a time, rinse worms 6 times with 1x Montjuic water to trigger release of mucus by the worms.
c. Exchange 1x Montjuic water for 10 mL 5% N-Acetyl Cysteine in 1x PBS.

i. Gently swirl the dish, ensuring animals are submerged and floating in solution. Set at timer for 6 minutes.
ii. Periodically give the dish a gentle swirl to ensure the animals don’t settle to the bottom. Be ready to remove the solution promptly when the time goes off.
d. Remove 5% N-Acetyl Cysteine promptly, keeping the animals corralled and submerged by tilting the plate and preventing exposure to air.
22. Add 10 mL 4% formaldehyde in 1x PBSTx, gently swirling dish to ensure worms do not adhere to the bottom of the dish or to one another. Set a timer for 1 hour.

a. Periodically swirl the dish during the fixation to ensure the animals don’t settle and stick to the bottom. Separate worms that associate with one another through prompt, gentle pipetting. **CRITICAL:** Perform formaldehyde fixation in a chemical fume hood while wearing PPE (lab coat, gloves, safety glasses). Follow hazardous waste collection and disposal guidelines for formaldehyde-containing solutions.
24. Remove the fixation solution and replace it with 10 mL 1x PBSTx, keeping the animals corralled and submerged by tilting the plate and preventing exposure to air.

a. Perform 3 x 5-minute washes with 1x PBSTx, squirting apart any worms that stick to one another.
b. Transfer worms to labeled 1.5 mL tubes after second wash (date, fragment type, dpc, target gene, dose, n) using a cut p200 tip.
25. Remove 1x PBSTx and replace with 1:1 v/v methanol:1x PBSTx for at least 5 minutes at room temperature.
26. Remove 1:1 v/v methanol:1x PBSTx and replace with 100% methanol for 5 minutes at room temperature.

a. Repeat 100% methanol exchange.
b. Store samples in 100% methanol at −20°C for at least one hour or long-term storage.

**CRITICAL:** Follow hazardous waste collection and disposal guidelines for methanol-containing solutions.

Note: Perform whole mount *in situ* hybridization on fixed fragments following protocols described in ^1,28,29^.

## Key resources table

| REAGENT or RESOURCE | SOURCE | IDENTIFIER |
| --- | --- | --- |
| **Antibodies** |  |  |
| N/A |  |  |
| **Chemicals, peptides, and recombinant proteins** |  |  |
| gentamicin sulfate powder | Gemini Bioproducts | #400-100 P |
| sodium chloride | EMD Millipore Sigma | Sigma # S3014 |
| sodium bicarbonate | EMD Millipore Sigma | Sigma # S5761 |
| potassium chloride | EMD Millipore Sigma | Sigma # P5405 |
| magnesium sulfate | EMD Millipore Sigma | Sigma # M2643 |
| calcium chloride dihydrate | EMD Millipore Sigma | Sigma # C7902 |
| dextrose (D+ glucose) | EMD Millipore Sigma | Sigma # G7021 |
| sodium phosphate, dibasic heptahydrate | EMD Millipore Sigma | Sigma # 2429 |
| potassium phosphate, monobasic | EMD Millipore Sigma | Sigma #5655 |
| Triton X-100 | EMD Millipore Sigma | Sigma #T8787 |
| N-acetyl L-cysteine | EMD Millipore Sigma | Sigma #A7250 |
| 37% formaldehyde aqueous solution | EMD Millipore Sigma | Sigma #F8775 |
| Clorox germicidal bleach | Clorox professional products company | B016OKJD9S |
| agarose (low-melting) | Fisher Scientific | BP165-25 |
| DAPI nucleic acid stain (10 mg) | ThermoFisher Scientific | D1306 |
| Invitrogen TRIzol reagent | ThermoFisher Scientific | 15596026 |
| chloroform | EMD Millipore Sigma | C2432 |
| Isopropanol, 99.5% for molecular biology, DNase, RNase, and Protease free | ThermoFisher Scientific | 327272500 |
| Ethanol absolute (200 proof), molecular biology grade | Fisher Scientific | BP2818100 |
| methanol | Fisher Scientific | A454-1 |
| **Critical commercial assays** |  |  |

| Deposited data |  |  |
| --- | --- | --- |
| *S. polychroa* transcriptome (dd_Spol_v4) | Planmine<sup>30</sup> | https://planmine.mpinat.mpg.de/planmine/begin.do |
| Experimental models: Organisms/strains |  |  |
| *S. polychroa* (3n=12) hermaphrodites | 1 | N/A |
| Oligonucleotides |  |  |
| Primers for cloning *Spol-β-catenin-1* dsRNA 1 and 2 | 1 | N/A |
| Primers for cloning *Spol-ovo* dsRNAs, *Spol-foxA* dsRNAs, and *GFP* dsRNA, see **Table 1** | This paper | N/A |
| Primers for qRT-PCR (*Spol-β-catenin-1*, *Spol-β-catenin-2*, *Spol-EF-1*) | 1,31 | N/A |
| Primers for qRT-PCR (*Spol-ovo*) **Table 2** | This paper | N/A |
| Recombinant DNA |  |  |
| pDL1-Spol-β-catenin-1dsRNA1 plasmid | 1 |  |
| pDL1-Spol-β-catenin-1 dsRNA2 plasmid | 1 |  |
| pDL1-Spol-ovo-N-term dsRNA plasmid | This paper | Addgene #242942 |
| pDL1-Spol-ovo-C-term dsRNA plasmid | This paper | Addgene #242943 |
| pDL1-Spol-foxA1-C-term dsRNA plasmid | This paper | Addgene #242944 |
| pDL1- -GFP dsRNA plasmid | This paper | Addgene #242945 |
| Other |  |  |
| Disposable PES bottle top filters (0.2 μm) | Fisher Scientific | FB12566510 |
| Samco™ Transfer Pipettes (691, non-sterile) | Thermo Fisher Scientific | 691TSPK |
| 35 x 10 mm Greiner Bio-One 627102 polystyrene vented dishes | Fisher Scientific | 07-000-327 |
| 100 mm x 15 mm polystyrene petri dishes | Fisher Scientific | FB087579B |
| dumont #5 fine forceps | Fine Science Tools | 11254-20 |
| insect pins (size 000) | Fine Science Tools | 26001-25 |
| moria nickel plated pin holder | Fine Science Tools | 26016-12 |
| micro knives (plastic handle, 22.5°) | Fine Science Tools | 10316-14 |

## Materials and equipment setup

**50 mg/mL gentamicin sulfate**

Materials and equipment setup **50 mg/mL gentamicin sulfate**
| Reagent | Final concentration | Amount |
| --- | --- | --- |
| Gentamicin sulfate | 50 mg/mL | 10 g |
| ddH<sub>2</sub>O | n/a | to 200 mL |
| **Total** | **n/a** | **200 mL** |

**5x Holfreter’s media pH 7.5, for dsRNA soaks**

**5x Holfreter's media pH 7.5, for dsRNA soaks**
| Reagent | Final concentration (5x) | Amount |
| --- | --- | --- |
| sodium chloride | 299.5 mM | 3.5 g |
| sodium bicarbonate | 11.9 mM | 0.2 g |
| potassium chloride | 3.35 mM | 0.05 g |
| magnesium sulfate | 8.3 mM | 0.2 g |
| calcium chloride dihydrate | 3.35 mM | 0.1 g |
| dextrose (D+ glucose) | 27.75 mM | 1.0 g |
| ddH<sub>2</sub>O | n/a | to 200 mL |
| **Total** | **n/a** | **200 mL** |

**1x Holfreter’s media pH 7.5 for planaria embryo culture**

**1x Holfreter's media pH 7.5 for planaria embryo culture**
| Reagent | Final concentration (1x) | Amount |
| --- | --- | --- |
| sodium chloride | 59.9 mM | 3.5 g |
| sodium bicarbonate | 2.38 mM | 0.2 g |
| potassium chloride | 0.67 mM | 0.05 g |
| magnesium sulfate | 1.66 mM | 0.2 g |
| calcium chloride dihydrate | 0.67 mM | 0.1 g |
| dextrose (D+ glucose) | 5.55 mM | 1.0 g |
| ddH<sub>2</sub>O | n/a | to 1L |
| **Total** | **n/a** | **1 L** |

Adjust to pH 7.5. Filter sterilize (0.2 µm membrane). Store at room temperature. Use within 2 months.

**10% bleach**

| Reagent | Final concentration | Amount |
| --- | --- | --- |
| clorox germicidal bleach | 10% | 10 mL |
| ddH<sub>2</sub>O | n/a | to 100 mL |
| **Total** | **n/a** | **100 mL** |

**CRITICAL:** Avoid contact between bleach and exposed skin, eyes, and clothing. Handle bleach-containing solutions with gloves, protective eyewear, and a lab coat.

**Optional reagents, needed for fixing samples at 14 dpc:**

**10x PBS**

| Reagent | Final concentration | Amount |
| --- | --- | --- |
| sodium chloride | 1.37 M | 80 g |
| potassium chloride | 27 mM | 2 g |
| sodium phosphate, dibasic heptahydrate | 101 mM | 27.2 g |
| potassium phosphate, monobasic | 18 mM | 2.4 g |
| **Total** | **n/a** | **1000 mL** |

Autoclave 20 minutes. Store at room temperature. Use within 1 year. pH of 1x PBS diluted from 10x PBS = 7.4.

**1x PBSTx (1x PBS + 0.5% Triton X-100)**

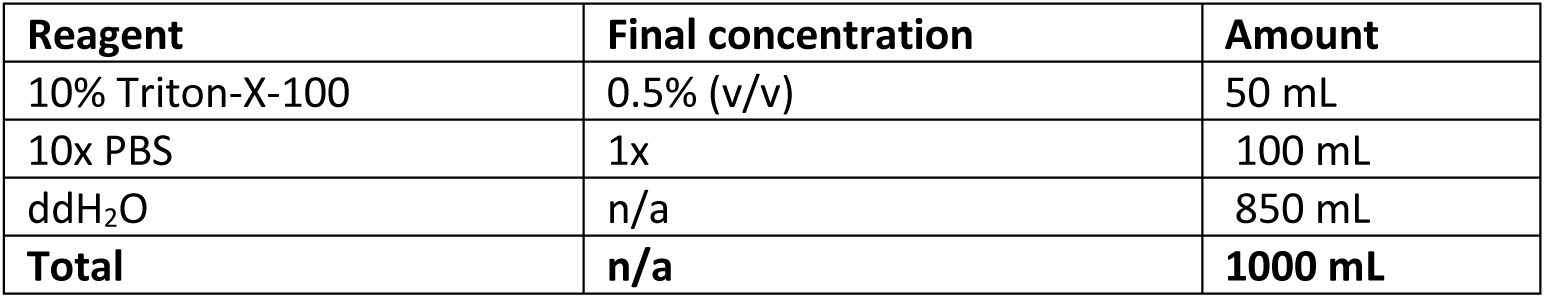

**5% N-acetyl cysteine**

| Reagent | Final concentration | Amount |
| --- | --- | --- |
| N-acetyl cysteine | 5% w/v | 0.5 g |
| 1x PBS | n/a | to 10 mL |
| **Total** | **n/a** | **10 mL** |

Make fresh with each use. Make 10 mL per sample (n~ 20 animals, per 60 mm dish).

**4% formaldehyde in 1x PBSTx**

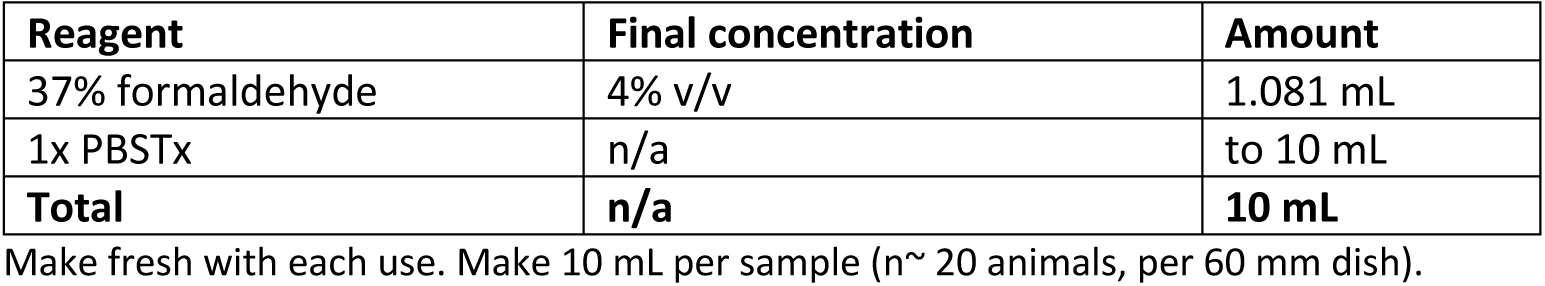

**CRITICAL:** Work in a chemical fume hood while wearing PPE (lab coat, gloves, safety glasses). Follow hazardous waste collection and disposal guidelines for formaldehyde-containing solutions.

**1:1 Methanol: 1x PBSTx**

| Reagent | Final concentration | Amount |
| --- | --- | --- |
| methanol | 50% (v/v) | 50 mL |
| 1x PBSTx | 50% (v/v) | 50 mL |
| **Total** | **n/a** | **100 mL** |

**CRITICAL:** Work in a chemical fume hood while wearing PPE (lab coat, gloves, safety glasses). Follow hazardous waste collection and disposal guidelines for methanol-containing solutions.

## Expected outcomes

We demonstrate efficacy of dsRNA soaking to produce penetrant RNAi knock-down phenotypes in regenerating *S. polychroa* S7 embryo fragments, expanding opportunities to interrogate gene function to understudied life cycle stages in planarian flatworms. We show proof-of-concept by targeting three genes that produce striking phenotypes visible in live specimens: *Spol-ß-catenin-1, Spol-ovo,* and *Spol-foxA.* We characterize knock-down phenotypes in fixed fragments using whole mount *in situ* hybridization (WISH) and demonstrate gene knock-down quantitatively by qRT-PCR.

In ^1^, we produced penetrant *Spol-ß-catenin-1* knock-down phenotypes using non-overlapping N-terminal and C-terminal dsRNAs (*ß-cat-1* dsRNA1 and dsRNA2, respectively). We determined the *Spol-ß-cat-1* dsRNA dose for these experiments empirically by incubating freshly cut S7 anterior and posterior fragments with 1, 10, 50, 100, 250 ng/µl *Spol-ß-cat-1* dsRNA for two days, and scored head regeneration phenotypes at 7 and 14 dpc. We determined 10 ng/µl *Spol-ß-cat-1* dsRNA 1 or 2 was sufficient to produce highly penetrant phenotypes in S7 anterior and posterior fragments. We did not vary the soak time post-cut, reasoning that access to internal tissues was greatest prior to wound closure. Ectopic head tissue produced by *Spol-ß-cat-1(RNAi)* was detectable by 7 dpc, but scoring was easier at 14 dpc, when regenerated ectopic eye(s) were larger.

Both *ß-cat-1* dsRNA 1 and 2 elicited polarity reversal phenotypes in S7 anterior fragments at high frequency, similar to the knock-down phenotype reported for adult asexual *S. mediterranea*^32–34^. Strikingly, *ß-cat-1(RNAi)* elicited head regeneration in S7 posterior fragments cut immediately anterior to the pharynx, which are normally unable to make new heads^1^. We performed WISH on fixed 14 dpc *Spol-ß-cat-1(RNAi)* fragments to corroborate the expansion of anterior territory and concomitant loss of posterior marker gene expression, and to visualize brain and eye tissue^1^. qRT-PCR assays were performed on *unc-22(RNAi)* negative control and *Spol-ß-cat-1(RNAi)* regenerates harvested at 14 dpc. A *Spol-ß-cat-1* qPCR amplicon, located between the sequences targeted by *Spol-ß-cat-1* dsRNA1 and 2, and an established housekeeping control, *Spol-EF-1*^31^, were used to determine the relative fold-change in gene expression between control and *Spol-ß-cat-1(RNAi)* samples using the 2^−ΔΔCt^ method^1^. Both *Spol-ß-cat-1* dsRNAs 1 and 2 significantly reduced expression of the intended target, but not the paralog *Spol-ß-cat-2,* demonstrating knock-down efficacy and specificity.

To further demonstrate the production of regeneration-defective phenotypes using this protocol, we performed RNAi knock-downs for *Spol-foxA* and *Spol-ovo,* two genes encoding fate specifying transcription factors whose asexual *S. mediterranea* homologs are required for the maintenance and regeneration of the pharynx and eyes, respectively^4,9,35^. In developing *S. polychroa* embryos and *S. mediterranea* asexual adults, *foxA* expression is restricted to mesenchymal pharynx-fated progenitors and differentiated pharynx tissues^1,9,35,36^. Similarly, *Smed-ovo* is expressed in eye progenitor cells and photoreceptor neurons and non-neuronal pigment cup cells in the eye^4^. We reasoned that we could effectively minimize pre-existing sources of expression for both targets through amputation. *Spol-foxA* knock-down experiments were performed using S7 anterior fragments (**Troubleshooting Problem 4, Figure 2B**), and *Spol-ovo* knock-downs were performed using S7 posterior fragments, cut within the head (**Troubleshooting Problem 5, Figure 3B**).

**Figure 2:**
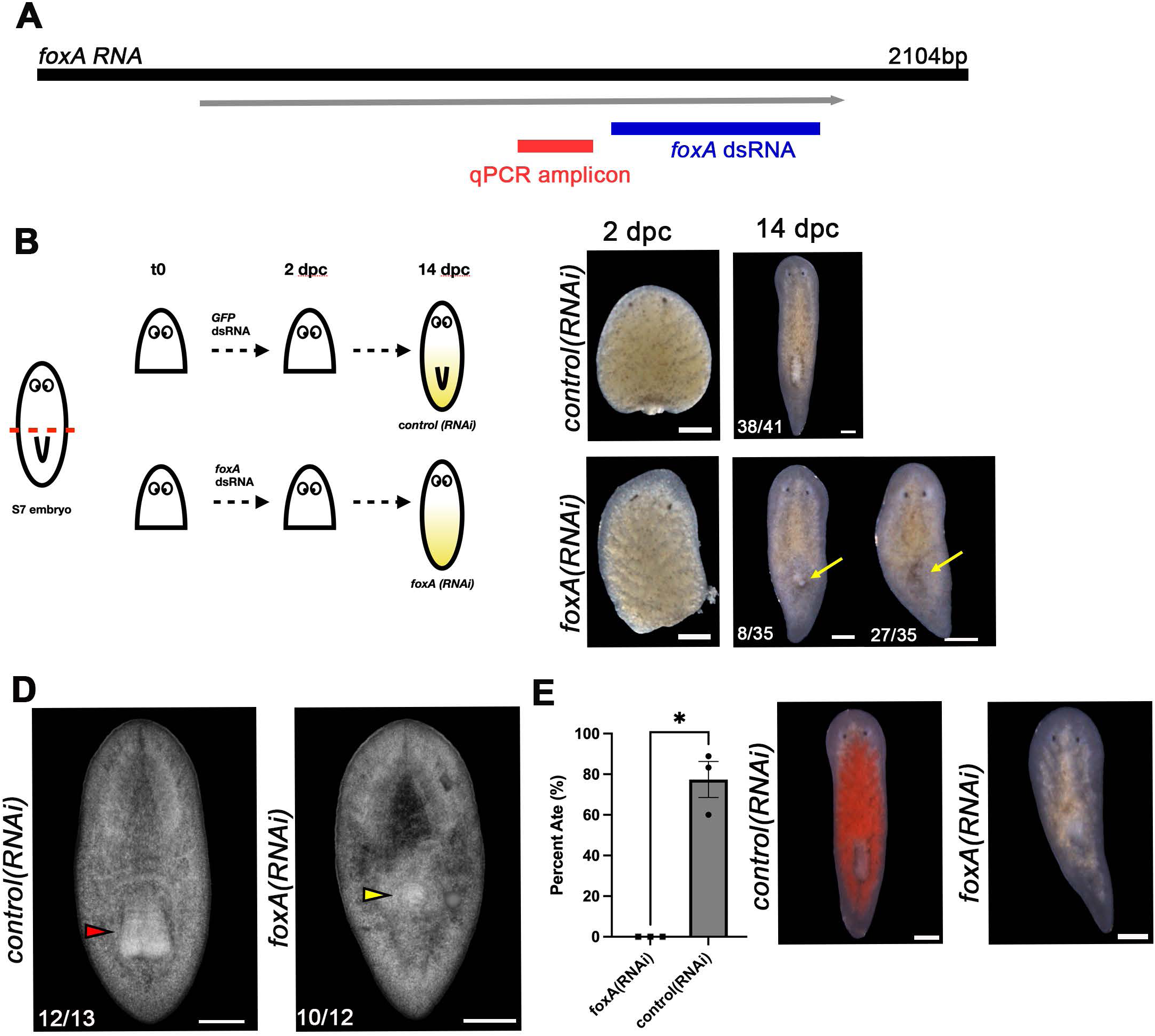
*Spol-fox-A* is required for definitive pharynx regeneration in S7 anterior fragments. **2A**. Schematic of the *Spol-foxA* coding sequence showing the coding sequence fragment cloned into pDL1 for dsRNA production and the qPCR amplicon. **2B**. S7 embryos were amputated immediately anterior to the pharynx to produce anterior fragments that were soaked in 250 ng/µl GFP dsRNA or *Spol-foxA* dsRNA for 48 hours, then moved to recovery media and cultured until 14 dpc, when they were scored visually, harvested for total RNA extraction, and challenged with a feeding assay. Regenerates were fixed at 20 dpc for DAPI staining. **2C**. Brightfield images of S7 anterior fragments treated with 250 ng/µl *GFP* dsRNA (*control(RNAi))* or *Spol-foxA* dsRNA (*foxA(RNAi)*) for 48 hours post-amputation, at 2 dpc and 14 dpc. Yellow arrow: Dorsal tissue outgrowths observed in regenerating *Spol-foxA(RNAi)* fragments. Anterior: Up. Scale bar: 200 µm. **2D**. DAPI staining on *control(RNAi)* and *Spol-foxA(RNAi)* S7 anterior fragments fixed at 20 dpc. Red arrowhead: normal pharynx morphology. Yellow arrowhead: failed pharynx regeneration. Anterior: Up. Dorsal views. Scale bar: 200 µm. **2E**. Left: Percent *control(RNAi)* and *Spol-foxA(RNAi)* S7 anterior fragment regenerates that ate liver after 1 hour feeding assay at 14 dpc. Right: *control(RNAi)* animals (left) show evidence of red liver throughout their gut, while *Spol-foxA(RNAi)* animals (right) did not ingest liver. Two-tailed paired T-test, p=0.0128. Scale bar: 200 µm. **2B-E**: 250 ng/µl dsRNA doses. 3 independent experiments. Error bars: standard error of the mean. *control(RNAi):* n=41. *Spol-foxA(RNAi)* n=35.

**Figure 3:**
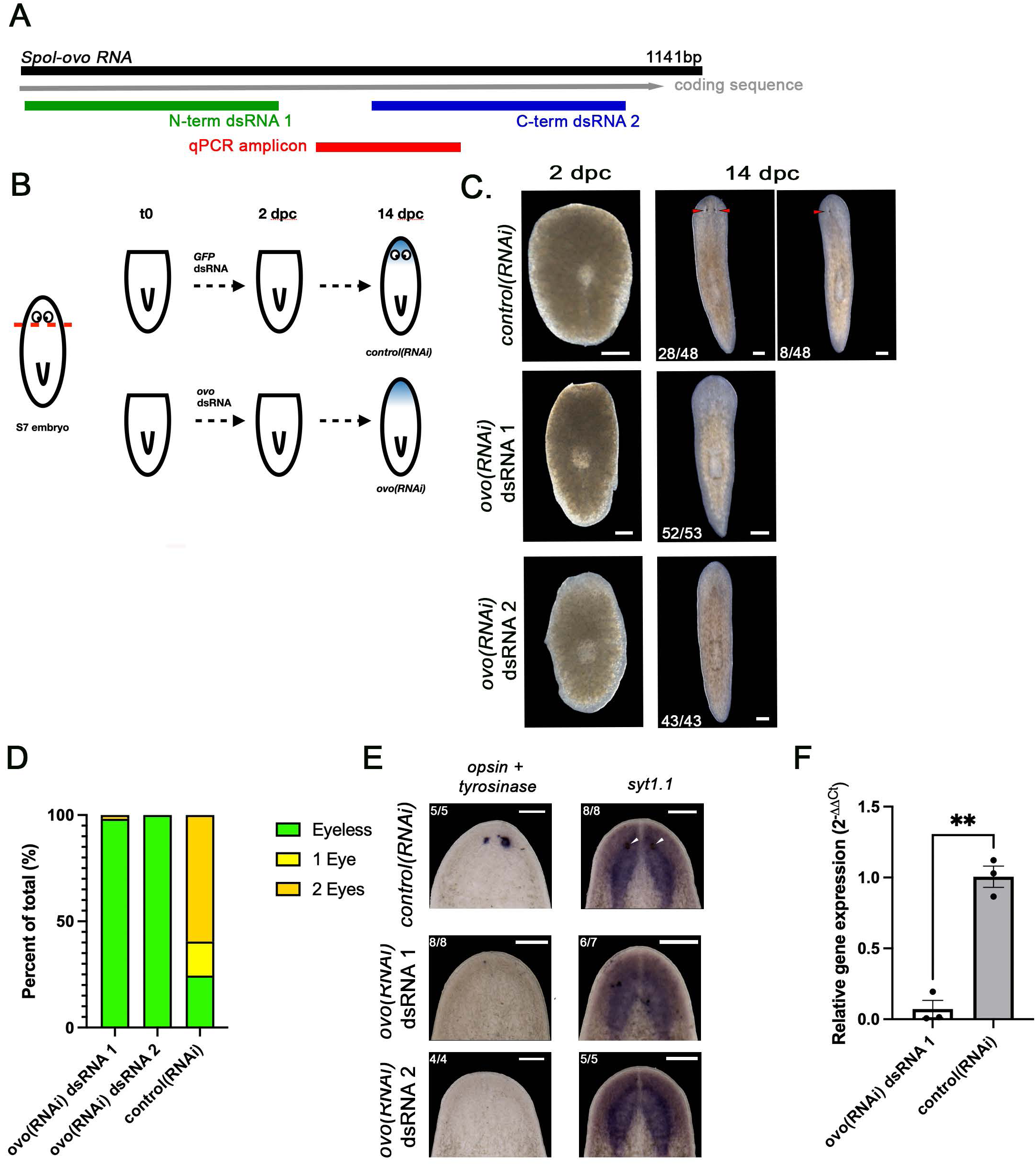
*Spol-ovo* is required for eye regeneration in S7 posterior fragments. **3A.** Schematic of the *Spol-ovo* coding sequence showing the N-terminal and C-terminal fragments cloned into pDL1 for dsRNA production and the locations of the qPCR amplicon. **3B**. S7 embryos were amputated beneath the eyes to produce posterior fragments that were soaked in 50 ng/µl GFP dsRNA or *Spol-ovo* dsRNA 1 or 2 for 48 hours, then moved to recovery media and cultured until 14 dpc, when they were scored visually, harvested for total RNA extraction, and fixed for whole mount in situ hybridization (WISH). **3C**. Brightfield images of S7 posterior fragment regenerates treated with 50ng/µl *GFP* dsRNA (*control(RNAi))* or *Spol-ovo* dsRNA 1 or 2 (*Spol-ovo(RNAi))*, at 2 dpc and 14 dpc. **3D.** Percent *control(RNAi)*, *Spol-ovo(RNAi)* dsRNA 1, and *Spol-ovo(RNAi)* dsRNA 2 S7 posterior fragments that regenerated a head with no eyes (green), 1 eye (yellow), or 2 eyes (orange) at 14 dpc. **3E**. WISH on *control(RNAi)* and *Spol-ovo(RNAi)* S7 posterior fragment regenerates fixed at 14 dpc, using *opsin + tyrosinase* riboprobes (regenerated eyes, left) or *syt1.1* (regenerated brain and ventral nerve cords). **3C, E**. Arrowheads: regenerated eyes. Lower left corner: number of animals exhibiting phenotype/total animals scored. Anterior: Up. Dorsal views. Scale bar: 200 µm. **3F**. qRT-PCR measurements of *Spol-ovo* expression in *(control)RNAi* and *Spol-ovo(RNAi)* S7 posterior fragment regenerates at 14 dpc. *Spol-ovo* dsRNA 1 treatment produced a significant decrease in expression relative to GFP dsRNA (Paired t test, two tailed, p=0.0093). **3B-E.** dsRNA dose: 50 ng/µl. 3 independent experiments. Error bars: standard error of the mean. *GFP(RNAi):* n=48. *Spol-ovo(RNAi)* dsRNA 1 n=53. *Spol-ovo(RNAi)* dsRNA 2 n=43.

We generated a plasmid DNA construct complementary to the *Spol-foxA* C-terminus (**Figure 2A**), and purified dsRNA from transformed HT115 bacteria as described in^26^ and **Troubleshooting Problem 2**. We determined that 48-hour soaks (**Troubleshooting Problem 3**) in 250 ng/µl *Spol-foxA* dsRNA were sufficient to produce fully penetrant pharynx regeneration phenotypes (**Figure 2B**). *control(RNAi*) S7 anterior fragments treated with 250 ng/µl GFP dsRNA regenerated the pharynx and tail tissue by 14 dpc (**Figure 2C-D**). In contrast, *Spol-foxA(RNAi)* S7 anterior fragments showed impaired pharynx regeneration while tail regeneration was unaffected (**Figure 2C-D**). In some instances, *Spol-foxA(RNAi)* S7 anterior fragments had small pharynx primordia at 7 dpc, while others appeared to have an empty pharyngeal pouch within the regenerated trunk (**Figure 2C-D**). Frequently, dark pigmentation marked the site of the pharynx pouch, and sometimes dorsal outgrowths were noted above the esophagus and pharyngeal pouch (**Figure 2C-D**). *Spol-foxA(RNAi)* phenocopied reported S*med-foxA1(RNAi)* phenotypes^9,35^. We performed feeding assays to vet regenerated pharynx function. At 14 dpc, *control(RNAi*) and *Spol-foxA(RNAi)* S7 anterior fragment regenerates were challenged to eat raw beef liver supplemented with red food coloring to visualize food uptake into the digestive system^9^. After 60 minutes, 77% *GFP(RNAi*) animals had eaten (**Figure 2E**). In contrast, none of the *Spol-foxA(RNAi)* ingested liver during the feeding assay (**Figure 2E**), consistent with failed pharynx regeneration.

We made plasmid DNA constructs for two non-overlapping dsRNAs complementary to the *Spol-ovo* N- and C-terminus (**Figure 3A** dsRNA 1 and dsRNA 2, respectively), and purified dsRNAs from transformed HT115 bacteria as described in^26^ and **Troubleshooting Problem 2**. We determined that 48-hour soaks in 50 ng/µl *Spol-ovo* dsRNA 1 or 2 produced fully penetrant eyeless phenotypes when S7 posterior fragments were amputated immediately posterior to the eyes (**Figure 3B, Troubleshooting Problem 3**). 75% of *control(RNAi*) S7 posterior fragments regenerated eyes by 14 dpc (**Figure 3C-D**) with 59% of animals regenerating both eyes (28/48) and 16% (8/48) regenerating 1 eye. Strikingly, *Spol-ovo(RNAi)* caused eye regeneration failure: 98% *ovo* dsRNA1 and 100% *ovo* dsRNA2-treated regenerates lacked eyes (**Figure 3C-D**), phenocopying results produced in *S. mediterranea* adults^4^. Eyeless phenotypes were confirmed by performing WISH on *control(RNAi*) and *Spol-ovo(RNAi)* 14 dpc regenerates using *Spol-opsin-1* and *Spol-tyrosinase* riboprobes, specific markers for the photoreceptor neurons and pigment cups, respectively (**Figure 3E**). To verify *Spol-ovo* knockdown did not impair regeneration of other structures in the head, we performed WISH for *Spol-syt1.1,* a neuronal marker, and observed robust regeneration of the brain in *control(RNAi*) and *Spol-ovo(RNAi)* heads(**Figure 3E**). Knock-down efficacy was confirmed in *Spol-ovo(RNAi)* animals treated with dsRNA1 relative to *control(RNAi)* at 14 dpc by qRT-PCR (**Figure 3F**).

### Limitations

This protocol requires access to an *S. polychroa* breeding colony capable of laying at least 100 gravid egg capsules per day and a husbandry regimen that includes daily egg capsule collections. A reliable supply of dated egg capsule collections is necessary to ensure that enough embryos at the desired developmental stage and size are collected to execute properly controlled experiments for multiple target genes and/or dsRNA doses. The reproductive output of embryo-producing planarians varies across strains and species, which may impact the ease and feasibility of protocol implementation. The design and cloning of dsRNA target sequences requires the existence of searchable transcriptomic resources, which may not be available for strains and/or species of interest. In such cases, designing degenerate primers using target gene sequence(s) from a closely related species may facilitate cloning and dsRNA production.

Successful implementation of this protocol is contingent on the ability to remove embryos from egg capsules without injuring them and to proper staging of embryos (**Figure 1A**, **Troubleshooting Problem 1**)^1,27^. Regenerative abilities of *S. polychroa* S7 fragments are dependent on cut plane position and wound site orientation^1^. Please refer to **Troubleshooting Problems 4 and 5** for additional information on cut paradigms and experimental design. We recommend performing transverse cuts on S7 embryos, since most S7 fragments successfully complete wound closure^1^. We have not performed dsRNA soaks on sagittal (midline) cuts, but we advise against performing sagittal cuts prior to S7.5 due to low fragment survival at earlier stages^1^. Efficacy of dsRNA soaks in amputated fragments may be impacted by the speed and efficacy of wound closure. Therefore, fragment soaking may be less efficacious at later developmental stages, when wound healing is quicker, and achieving knock-down may require dsRNA soaking at higher doses, increased soak time, or combining soaking with dsRNA injection (**Troubleshooting Problem 3**). We have not tested whether RNAi knock-downs can be achieved in intact *S. polychroa* embryos through dsRNA soaking, nor have we tried soaking intact juvenile or adult animals.

RNAi knock-downs achieve post-transcriptional gene silencing and are not stable gene knockouts. The extent and efficacy of knock-downs vary and depend on many factors, including the site(s) and level of target gene expression, transcript stability, as well as target protein levels and stability. Some genes may not be amenable to RNAi knock-down in the described assay system. In addition, some target genes may be knocked down effectively (assessed by qRT-PCR) but may not produce a phenotype. Finally, this dsRNA soaking protocol, like all planarian RNAi methods, cannot be deployed in a tissue-specific manner. Assigning requirements for gene function in particular tissue(s) of interest may be difficult using whole-animal RNAi phenotypes.

## Troubleshooting

### Problem 1: Staging S7 *S. polychroa* embryos

Embryo staging relies on chronological, morphological, and behavioral criteria that are interpreted along a developmental continuum. Staging decisions are inherently observation-based and subjective categorizations, introducing variability to the process.

#### Potential solution

- We provide detailed descriptions of salient staging criteria for S6-S7.5 (**Step 8d. i-iv**) and representative live images (**Figure 1c**).
- We recommend bleaching and cracking open several egg capsules from 2-3 consecutive collection dates to determine the cohort that contains the most embryos of the desired stage. The collection date is 1 day post-egg capsule deposition (dped). See **Figure 1A** and **Step 8d. i-iv.**
- Cracking open egg capsules and removing embryos requires practice. We suggest learning to dissect later developmental stages (S8, approximately 12-14 dped), when embryos are well formed and they can swim out of the shells on their own, before trying dissections at earlier stages.
- If egg capsule number or embryo count is limiting, contact the authors to discuss animal colony husbandry practices that may improve egg capsule fertility and fecundity.

### Problem 2: Maximizing dsRNA production using HT115 bacterial cultures

Some target genes may require high dsRNA doses to achieve penetrant knock-down phenotypes. Alternatively, researchers may wish to purify large quantities of dsRNA for use in several experiments.

#### Potential solution

- We recommend doubling the volume of HT115 bacterial culture to 40mL during the induction or inducing two HT115 cultures of the desired construct at 20mL^26^. This ensures enough dsRNA for multiple replicates if desired. Cell pellets can be stored at −20°C for a few months, if necessary, prior to purification.
- IPTG induction for 4-5 hours is sufficient to produce enough purified dsRNA for an experiment with multiple doses or several biological replicates.
- Though not required, making the lysis buffer fresh prior to extracting will help to maximize recovery of purified dsRNA at high concentration.

### Problem 3: Optimizing variables that may impact knock-down efficacy

Some genes may be easier to knock-down than others. The dsRNA dose required to achieve penetrant knock-down phenotypes may vary for different genes.

#### Potential solution

- First, we recommend testing experimental dsRNAs at a high dose (e.g., 500 ng/µl). If a penetrant phenotype is observed in high dose soaks, then we recommend assaying whether comparable results can be achieved with dsRNA soaks at lower doses (e.g., 10 ng/µl, 50 ng/µl, 100 ng/µl, 250 ng/µl, 500 ng/µl).
- Preincubating intact embryos in dsRNA-containing media, increasing soak duration, combining dsRNA soaking and injections, or combining two or more dsRNAs in soaks and/or injections may increase knock-down severity and penetrance.
- Genes that are lowly expressed prior to amputation may be easier to knock-down than genes that are highly expressed in intact embryos prior cutting.

### Problem 4: S7 anterior fragments: where should the cut plane be positioned?

Posterior (trunk and tail-forming) regeneration abilities are robust in S7 *S. polychroa* embryos but still show some variation with position of the cut plane along the anterior-posterior axis^1^.

#### Potential solution

We recommend making a single transverse cut immediately anterior to the developing pharynx to generate S7 anterior fragments for RNAi assays (**Figure 2B**). The vast majority of negative control-treated S7 anterior fragments cut at this position are expected to wound heal within 2 dpc and to regenerate missing posterior tissues within 7–14 dpc^1^. We advise against creating S7 anterior fragments from cuts within the head for three reasons: 1) regeneration incidence varied most at the most anterior cut position assayed in S7 embryos^1^, 2) S7 head amputations are difficult to standardize without visible morphological landmarks, and 3) head fragments will be small, likely less than 100-200µm in length, and will be difficult to handle.

### Problem 5: S7 posterior fragments: where should the cut plane be positioned?

Anterior (head-forming) regeneration abilities vary greatly depending on where S7 *S. polychroa* embryos are cut along the anteroposterior axis^1^. Amputations within the head produce posterior fragments that regenerate new head tissue at high frequency. In contrast, cuts made immediately anterior to the pharynx produce head regeneration-incompetent posterior fragments.

#### Potential solution

- To assay for regeneration of missing head structures (e.g., the brain, or eyes) we recommend amputating S7 embryos within the head (**Figure 3B**).
- To assay for precocious induction of head regeneration, we recommend cutting S7 animals immediately anterior to the pharynx. Please refer to *Spol-ß-cat-1* knock-down assays in ^1^.

## Resource availability

### Lead contact

Inquiries and requests for resources or reagents should be directed to and will be fulfilled by Erin Davies.

### Technical contact

Technical questions about protocol implementation should be addressed to Erin Davies.

### Materials availability

Plasmids generated for this study are available at Addgene (see Key Resources Table for details).

### Data and code availability

Neither data sets nor code were generated for this study.

## Acknowledgments

We thank Andrew Wolff and Daniel Lobo for discussions regarding the pDL1 Gibson assembly cloning pipeline, Stjin Mouton, Kirill Ustyantsev and Eugene Berezikov for sharing their protocol for dsRNA purification prior to publication, and members of the NCI-Frederick Cancer and Developmental Biology Laboratory for their constructive feedback and enthusiasm for our work. This research was supported by the Intramural Research Program of the National Institutes of Health (NIH) project number ZIA BC 0120009 (ELD). The contributions of the NIH author(s) were made as part of their official duties as NIH federal employees, are in compliance with agency policy requirements, and are considered Works of the United States Government. However, the findings and conclusions presented in this paper are those of the author(s) and do not necessarily reflect the views of the NIH or the U.S. Department of Health and Human Services.

## Author contributions

Conceptualization: EWD, CLTB, ELD. Methodology: EWD, CLTB, ELD. Writing – original draft: EWD, ELD. Writing – review and editing: EWD, CLTB, ELD. Supervision and funding acquisition: ELD.

---

**Table 1:**
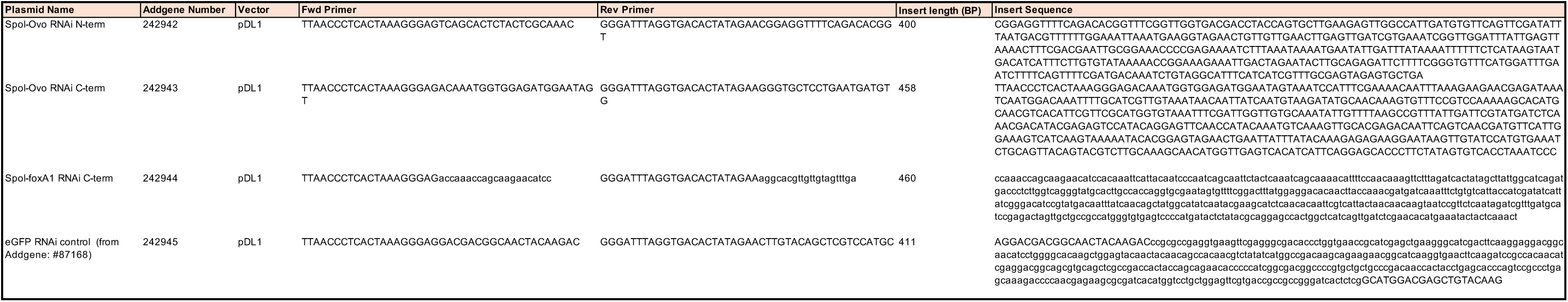
Primers and insert sequences for pDL1 constructs used for dsRNA production in this study.

**Table 2:**
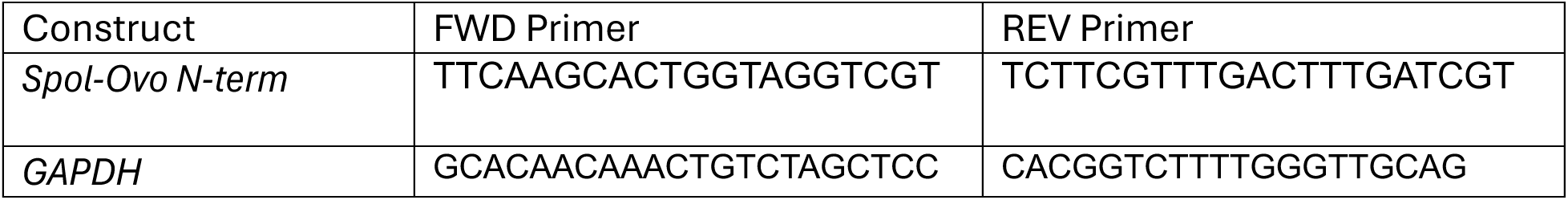
Primers for qRT-PCR assays to determine efficacy of *Spol-ovo* knock-downs.

## References

1. Booth, C.L.T., Stevens, B.C., Stubbert, C.A., Kallgren, N.T., Deihl, E.W., and Davies, E.L. (2025). Developmental onset of planarian whole-body regeneration depends on axis reset. Curr. Biol. 35, 2479–2494.

2. Wagner, D.E., Ho, J.J., and Reddien, P.W. (2012). Genetic regulators of a pluripotent adult stem cell system in planarians identified by RNAi and clonal analysis. Cell Stem Cell 10, 299–311.

3. Labbé, R.M., Irimia, M., Currie, K.W., Lin, A., Zhu, S.J., Brown, D.D.R., Ross, E.J., Voisin, V., Bader, G.D., Blencowe, B.J., et al. (2012). A Comparative Transcriptomic Analysis Reveals Conserved Features of Stem Cell Pluripotency in Planarians and Mammals. Stem Cells 30, 1734–1745.

4. Lapan, S.W., and Reddien, P.W. (2012). Transcriptome Analysis of the Planarian Eye Identifies *ovo* as a Specific Regulator of Eye Regeneration. Cell Rep. 2, 294–307.

5. Forsthoefel, D.J., James, N.P., Escobar, D.J., Stary, J.M., Vieira, A.P., Waters, F.A., and Newmark, P.A. (2012). An RNAi screen reveals intestinal regulators of branching morphogenesis, differentiation, and stem cell proliferation in planarians. Dev. Cell 23, 691–704.

6. Cowles, M.W., Brown, D.D.R., Nisperos, S.V., Stanley, B.N., Pearson, B.J., and Zayas, R.M. (2013). Genome-wide analysis of the bHLH gene family in planarians identifies factors required for adult neurogenesis and neuronal regeneration. Development 140, 4691–4702.

7. King, H.O., Owusu-Boaitey, K.E., Fincher, C.T., and Reddien, P.W. (2024). A transcription factor atlas of stem cell fate in planarians. Cell Rep. 43, 113843.

8. Scimone, M.L., Cote, L.E., and Reddien, P.W. (2017). Orthogonal muscle fibres have different instructive roles in planarian regeneration. Nature 551, 623–628.

9. Adler, C.E., Seidel, C.W., McKinney, S.A., and Sánchez Alvarado, A. (2014). Selective amputation of the pharynx identifies a FoxA-dependent regeneration program in planaria. Elife 3, e02238.

10. Khan, U.W., and Newmark, P.A. (2022). Somatic regulation of female germ cell regeneration and development in planarians. Cell Rep. 38, 110525.

11. Arnold, C.P., Benham-Pyle, B.W., Lange, J.J., Wood, C.J., and Sánchez Alvarado, A. (2019). Wnt and TGFβ coordinate growth and patterning to regulate size-dependent behaviour. Nature 572, 655–659.

12. Rouhana, L., Tasaki, J., Saberi, A., and Newmark, P.A. (2017). Genetic dissection of the planarian reproductive system through characterization of *Schmidtea mediterranea* CPEB homologs. Dev. Biol. 426, 43–55.

13. Reddien, P.W., Bermange, A.L., Murfitt, K.J., Jennings, J.R., and Sánchez Alvarado, A. (2005). Identification of genes needed for regeneration, stem cell function, and tissue homeostasis by systematic gene perturbation in planaria. Dev. Cell 8, 635–649.

14. Benham-Pyle, B.W., Brewster, C.E., Kent, A.M., Mann, F.G., Jr, Chen, S., Scott, A.R., Box, A.C., and Sánchez Alvarado, A. (2021). Identification of rare, transient post-mitotic cell states that are induced by injury and required for whole-body regeneration in *Schmidtea mediterranea*. Nat. Cell Biol. 23, 939–952.

15. Roberts-Galbraith, R.H., Brubacher, J.L., and Newmark, P.A. (2016). A functional genomics screen in planarians reveals regulators of whole-brain regeneration. Elife 5, e17002.

16. Sánchez Alvarado, A., and Newmark, P.A. (1999). Double-stranded RNA specifically disrupts gene expression during planarian regeneration. Proc. Natl. Acad. Sci. U. S. A. 96, 5049–5054.

17. Adler, C.E., and Alvarado, A.S. (2018). Systemic RNA interference in planarians by feeding of dsRNA containing bacteria. Methods Mol. Biol. 1774, 445–454.

18. Shibata, N., and Agata, K. (2018). RNA interference in planarians: Feeding and injection of synthetic dsRNA. Methods Mol. Biol. 1774, 455–466.

19. Orii, H., Mochii, M., and Watanabe, K. (2003). A simple “soaking method” for RNA interference in the planarian *Dugesia japonica*. Dev. Genes Evol. 213, 138–141.

20. Martín-Durán, J.M., Duocastella, M., Serra, P., and Romero, R. (2008). New method to deliver exogenous material into developing planarian embryos. J. Exp. Zool. B Mol. Dev. Evol. 310, 668–681.

21. Cebrià, F., and Newmark, P.A. (2005). Planarian homologs of netrin and netrin receptor are required for proper regeneration of the central nervous system and the maintenance of nervous system architecture. Development 132, 3691–3703.

22. Newmark, P.A., Reddien, P.W., Cebrià, F., and Sánchez Alvarado, A. (2003). Ingestion of bacterially expressed double-stranded RNA inhibits gene expression in planarians. Proc. Natl. Acad. Sci. U. S. A. 100 Suppl 1, 11861–11865.

23. Collins, J.J., 3rd, Hou, X., Romanova, E.V., Lambrus, B.G., Miller, C.M., Saberi, A., Sweedler, J.V., and Newmark, P.A. (2010). Genome-wide analyses reveal a role for peptide hormones in planarian germline development. PLoS Biol. 8, e1000509.

24. Wolff, A., Wagner, C., Wolf, J., and Lobo, D. (2022). In situ probe and inhibitory RNA synthesis using streamlined gene cloning with Gibson assembly. STAR Protoc 3, 101458.

25. Rouhana, L., Weiss, J.A., Forsthoefel, D.J., Lee, H., King, R.S., Inoue, T., Shibata, N., Agata, K., and Newmark, P.A. (2013). RNA interference by feeding in vitro-synthesized double-stranded RNA to planarians: methodology and dynamics. Dev. Dyn. 242, 718–730.

26. Mouton, S., Mougel, A., Ustyantsev, K., Dissous, C., Melnyk, O., Berezikov, E., and Vicogne, J. (2024). Optimized protocols for RNA interference in *Macrostomum lignano*. G3 (Bethesda) 14. 10.1093/g3journal/jkae037.

27. Cardona, A., Hartenstein, V., and Romero, R. (2005). The embryonic development of the triclad *Schmidtea polychroa*. Dev. Genes Evol. 215, 109–131.

28. King, R.S., and Newmark, P.A. (2013). In situ hybridization protocol for enhanced detection of gene expression in the planarian *Schmidtea mediterranea*. BMC Dev. Biol. 13, 8.

29. Gaetano, A.J., and King, R.S. (2023). A simplified and rapid in situ hybridization protocol for planarians. Biotechniques 75, 231–239.

30. Rozanski, A., Moon, H., Brandl, H., Martín-Durán, J.M., Grohme, M.A., Hüttner, K., Bartscherer, K., Henry, I., and Rink, J.C. (2019). PlanMine 3.0-improvements to a mineable resource of flatworm biology and biodiversity. Nucleic Acids Res. 47, D812–D820.

31. Sureda-Gómez, M., Martín-Durán, J.M., and Adell, T. (2016). Localization of planarian β-CATENIN-1 reveals multiple roles during anterior-posterior regeneration and organogenesis. Development 143, 4149–4160.

32. Gurley, K.A., Rink, J.C., and Sánchez Alvarado, A. (2008). Beta-catenin defines head versus tail identity during planarian regeneration and homeostasis. Science 319, 323–327.

33. Petersen, C.P., and Reddien, P.W. (2008). *Smed-betacatenin-1* is required for anteroposterior blastema polarity in planarian regeneration. Science 319, 327–330.

34. Iglesias, M., Gomez-Skarmeta, J.L., Saló, E., and Adell, T. (2008). Silencing of *Smed-betacatenin-1* generates radial-like hypercephalized planarians. Development 135, 1215–1221.

35. Scimone, M.L., Kravarik, K.M., Lapan, S.W., and Reddien, P.W. (2014). Neoblast specialization in regeneration of the planarian *Schmidtea mediterranea*. Stem Cell Reports 3, 339–352.

36. Martín-Durán, J.M., Amaya, E., and Romero, R. (2010). Germ layer specification and axial patterning in the embryonic development of the freshwater planarian *Schmidtea polychroa*. Dev. Biol. 340, 145–158.

